# A modified Rayleigh-Plesset equation for a liquid-crystalline shelled microbubble

**DOI:** 10.1101/606632

**Authors:** James Cowley, Anthony J. Mulholland, Anthony Gachagan

## Abstract

Premanufactured shelled microbubbles composed of a protein shell are currently licensed as ultrasound imaging contrast agents. Current research is focussing on using the protein shelled microbubbles as transportation mechanisms for localised drug delivery particularly in the treatment of various types of cancer. For the very first time, a theoretical model is developed for an incompressible, gas loaded shelled microbubble with a thin shell composed of a liquid-crystalline material. We show that liquid-crystalline shelled microbubbles exhibit significantly different physical characteristics from commercial protein shelled microbubbles such as Sonovue and Optison. The authors propose that these significantly different physical characteristics may enhance localised drug delivery. We use the technique of linearisation to predict the shelled microbubble’s natural frequency and relaxation time. These physical parameters strongly influence sonoporation which is the mechanism that is used for localised drug delivery. The influence of the material properties of the shell on the natural frequency and relaxation time are discussed. We have discovered that liquid-crystalline shelled microbubbles have a relaxation time that is 10 times longer than Sonovue and Optison.

## 1 Introduction

Premanufactured shelled microbubbles are currently licensed in the UK as ultrasound imaging contrast agents. Current research is focussing on using the microbubbles as a transportation mechanism for localised drug delivery specifically in the treatment of various cancers [1–9]. Ultrasound contrast agents (UCAs) are shelled microbubbles typically composed of a layer or several layers of a protein shell encapsulating a perfluoro gas that helps to stabilise the microbubble when it is injected into the bloodstream [10–12]. The shelled microbubbles have a typical radius of between 1 and 4 *μ*m allowing them to propagate through the capillaries in the human body and a shell thickness that varies between 4 and 100 nm depending on whether the UCA is a monolipid or polymer variant [13]. UCAs create a contrast with the surrounding tissue primarily due to an impedance mismatch with the surrounding fluid and the production of higher harmonics. Microbubbles resonate with typical frequencies in the range of 1 to 10 MHz producing nonlinear, multiple harmonic signals that enhance the quality of the medical imaging process [14]. There has been a research momentum growing in recent years to use the UCAs as localised drug delivery agents [15]. Much progress has been made but much remains to be done before this can be deployed routinely in patients [16]. Hence there is a need to develop virtual simulation tools to better understand the challenges. This paper contributes to this effort by identifying an entirely new type of shelled microbubble that is composed of a nematic liquid crystal, and discusses how the material parameters of the liquid-crystalline shell influences the dynamics of the shelled microbubble. Note that all the previous published literature pertaining to the modelling of shelled microbubbles focusses solely on protein shells. We show that nematic liquid-crystalline shells display significantly different physical characteristics from conventional protein shells: these physical characteristics are highly advantageous to the mechanism of sonoporation [17].

Most current shelled microbubble models are based on the Rayleigh-Plesset equation for a free gas bubble, which is derived by applying pressure balances to the inner surface of the shelled microbubble with those acting on the outside of the shelled microbubble’s surface and the surrounding liquid [18–20]. The Rayleigh-Plesset equation assumes that the microbubble oscillations are purely radial and that the surrounding liquid is incompressible. The gas in the shelled microbubble is assumed to behave adiabatically despite its polytropic index being relatively close to one which is associated with isothermal behaviour [20]. Whilst it is not fully modelled, most of these equations handle to some degree fluid compressibility. Viscous damping associated with the microbubble shell is also modelled in these equations.

Thin monolipid microbubbles have shells that are viscoelastic in nature, and behave more like a fluid than a solid shell [20]. This fluid like behaviour has inspired us to consider one particular type of mesophase material, specifically liquid crystals. We propose a new type of shelled microbubble that is composed of a thin liquid-crystalline shell. This paper uses the Leslie-Erikson continuum theory ([21], p133-159) for liquid crystals to build up a model for the dynamics of the shelled microbubble. This is the first study that has used liquid crystal theory to model UCAs. This paper also considers, for the first time, how both the relaxation time which is defined as the time taken for the amplitude of an oscillating shelled microbubble to decrease to 1/*e* of its original amplitude, and the natural frequency of the microbubble are influenced by the material parameters of the shell such as the shell’s viscosity, density, thickness and its surface tension. The paper is structured in the following way: Section 2 deals with the generic Rayleigh-Plesset equation then Section 3 focusses on the evaluation of the stress of the liquid crystal’s shell with Section 4 considering the elastic energy density of the shell. Section 5 determines the linearised Rayleigh-Plesset model and Section 6 reports the results.

## 2 The Rayleigh-Plesset model

Consider a shelled microbubble with inner and outer radii given by *R*_1_ and *R*_2_ respectively, where the radii are functions of time only and the density of the shell is denoted by *ρ_S_*. This article uses a dot notation above a physical quantity to represent differentiation of that quantity with respect to time. In terms of tensor notation, let *x*_*i*_ represent the positional coordinate and *r* = |**x**| where *r*^2^ = *x*_*i*_*x*_*i*_. We shall denote the radial unit vector as *e*_*r*_ and the speed and acceleration of the inner radius of the microbubble as *Ṙ*_1_ and *R̈*_1_ respectively. Let *ρ*_*L*_ denote the density of the surrounding incompressible liquid where *σ* represents the Cauchy stress. Momentum balance results in the following equation to describe the dynamics of the UCA [18,22,23]

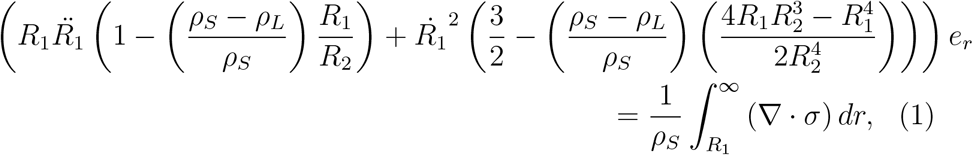

where a pressure balance has to be applied in order to determine the right hand side of equation (1). The pressure of the gas phase inside the shell and the surrounding ambient fluid pressure have to be considered as do the surface tensions and the shell and fluid viscosities. The divergence of the stress *σ* can be expressed as

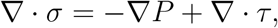

where *P* denotes a pressure term and *τ* represents both the stress in the shell and the stress due to the surrounding Newtonian fluid. Rewriting the right handside of equation (1) and integrating over the various media leads to

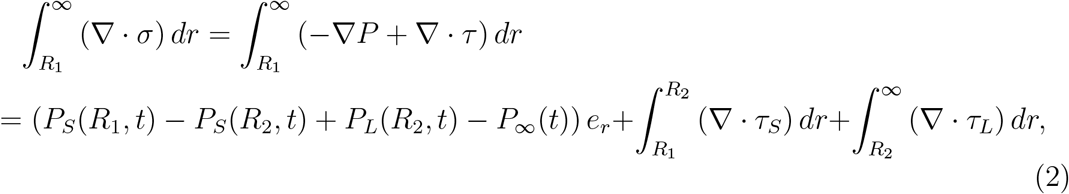

where *P*_*S*_, *P*_*L*_ and *P*_*∞*_ are the pressures in the shell, the surrounding Newtonian fluid, and at infinity, respectively. The stresses in the shell and the stress associated with the viscosity of the surrounding fluid are denoted by *τ*_*S*_ and *τ*_*L*_ respectively. Let *R*_01_ and *R*_02_ denote the equilibrium (unperturbed) inner and outer radius of the shelled microbubble. The boundary conditions at the inner and outer radii of the shell’s surface respectively are found by applying the momentum balance law [18] which leads to

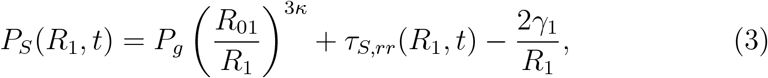

and

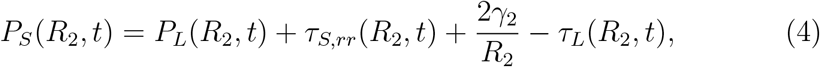

where *κ* denotes the polytropic index which is a dimensionless parameter [18,24] and *τ*_*S,rr*_ denotes the stress in the radial direction. The terms *γ*_1_ and *γ*_2_ denote the interfacial surface tension (gas-shell interface) and the surface tension between the outer shell and the surrounding liquid respectively. The gas pressure *P*_*g*_ in equation (3) is obtained by balancing the pressures at the equilibrium radii *R*_01_ and *R*_02_ to give

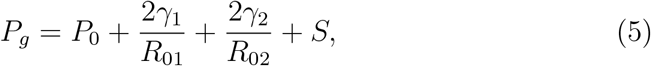

where *P*_0_ represents the surrounding ambient liquid pressure and *S* is the stress associated with the elastic energy density of the liquid crystal and is given by equation (23) in Section 4. Note that *P*_*∞*_ in equation (2) describes the atmospheric pressure plus any external applied pressures (such as those created by an ultrasound probe) and is represented by *P*_*∞*_ = *P*_0_ + *P*_A_ sin *ωt* where *P*_*A*_ and *ω* represent the externally applied pressure and angular frequency respectively. Substituting equations (3) and (4) into equation (2) gives

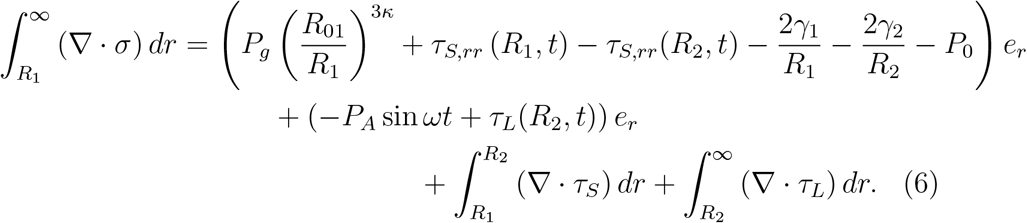

The stress due to the viscosity *μ*_*L*_ of the surrounding Newtonian fluid is denoted by *τ*_*L*_(*R*_2_, *t*) where ([18], [25] p50)

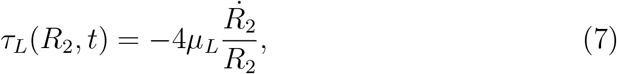

whereas the term 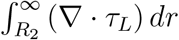 in equation (6) ([26], p354-p355) results in

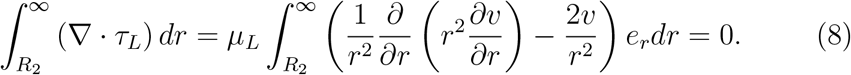

## 3 Calculating the stress of a liquid crystal shell

This section focusses on deriving an expression for the viscous stress of an incompressible liquid-crystal shell of known inner and outer radii. It is assumed that the shell’s composition is a liquid crystal that can be described dynamically using the nematic theory developed by Leslie and Ericksen ([21], p133-159) where five independent Leslie viscosities [27] are required to determine the stress in the shell. The sixth Leslie viscosity can be written as a linear combination of some of the other five independent Leslie viscosities. Some proteins [28] exhibit the characteristic behaviour of a liquid crystal where the molecules are arranged in layers ([21], p6). This paper will use nematic theory to model the mesophase behaviour of proteins [29]. Continuum modelling of liquid-crystal theory assumes that the molecules are rod like in nature and are described by a unit vector *n* which is called the director. The molecules are arranged in layers with the director aligning perpendicular to the layers and parallel to the layer normal ([21],p6). We shall assume spherical symmetry of the liquid-crystalline shell with the director pointing radially outward everywhere and the layers consisting of concentric spheres. The director describes the local direction of the average molecular alignment and is a unit vector (so *n* = *e*_*i*_*x*_*i*_/*r*) ([21],p6), where *x*_*i*_ represents the positional coordinate and *r* = |**x**|. The viscous stress *τ*_*ij*_ for a nematic liquid-crystal is given by

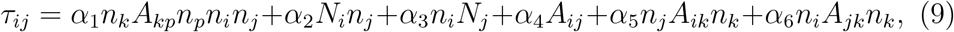

where *α*_1_, *α*_2_,…., *α*_6_ are the Leslie viscosities, *A*_*ij*_ is the rate of strain tensor and *N*_*i*_ is the co-rotational time flux of the director *n*. The co-rotational time flux is a measure of the rotation of the director, *n*, relative to the fluid. These terms are explicitly defined as *n*_*i*_ = *x*_*i*_/*r*, *A*_*ik*_ = (*v*_*i,k*_ + *v*_*k,i*_)/2, *N*_*i*_ = *ṅ*_*i*_ − *W*_*ij*_*n*_*j*_ where the superposed dot signifies the material time derivative *ṅ*_*i*_ = *∂n*_*i*_/*∂t* + *v*_*j*_*∂n*_*i*_/*∂x*_*j*_ and *W*_*ij*_ = (*v*_*i,j*_ − *v*_*j,i*_)/2 is the vorticity tensor. For the spherically symmetric case we have a velocity profile given by **v**= *ve*_*r*_ which is rewritten as

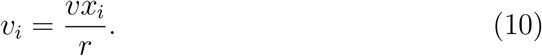

Hence

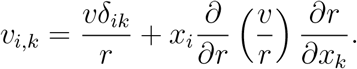

Since *r*^2^ = *x*_*k*_*x*_*k*_ then

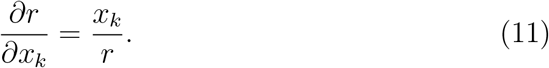

and so

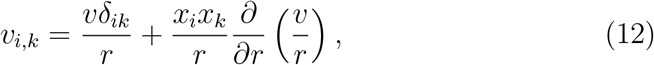

and since *δ*_*kp*_ = *δ*_*pk*_ then

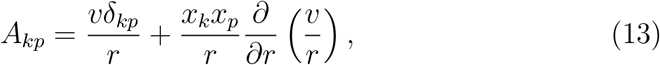

with *N*_*i*_ = 0 and *W*_*i,j*_ = 0. Substituting into equation (9) gives

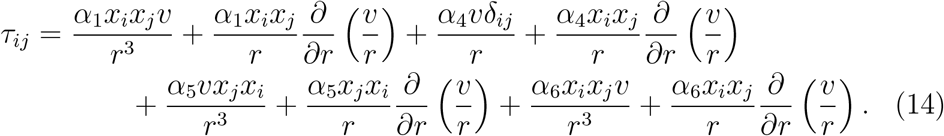

The shelled microbubble is assumed to be an incompressible shell composed of a thin liquid crystal shell with a radially directed flow ([21],p139). Marmottant et al. [19] discusses the limitations of an incompressible shell but only in relation to protein shells. UCAs such as Sonovue exhibit compression only behaviour [19]. Currently we have no experimental evidence as to whether or not compression only behaviour occurs for nematic liquid-crystalline shells. Since the shell is incompressible then its volume, *V*, and density will be time independent. For a shelled microbubble with an inner and outer radii given by *R*_1_ and *R*_2_ respectively, the following relationship holds

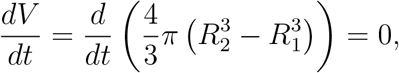

from which we can deduce that

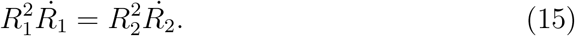

Church [18] and Doinikov et al. [22] state that

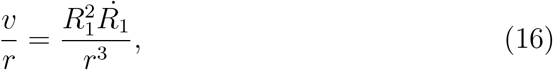

and so

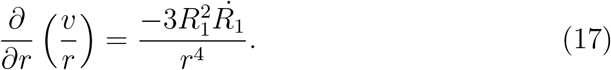

Using equations (16) and (17), the Leslie viscosities represented by equation (14) can be rewritten as

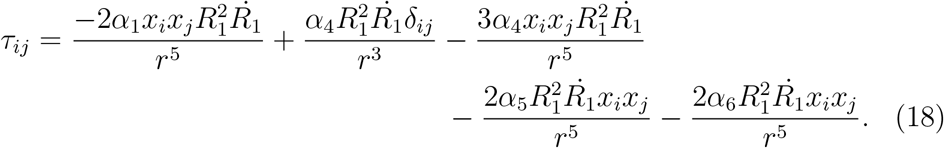

To determine the Cauchy momentum represented by equation (1) we have to evaluate the divergence of equation (18). Writing this in component form results in

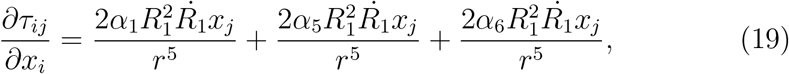

where the *α*_4_ contibution is zero which is consistent with Brennan ([25], p49-50). The stress associated with the Leslie viscosities is calculated by integrating equation (19) between the inner and outer radius of the shell. Our mathematical model focusses on purely radial oscillatory behaviour. Since the shelled microbubble moves solely in the radial direction then in spherical polar coordinates **r** = *re*_*r*_, with *r*^2^ = *x*_*j*_*x*_*j*_. The *j*th Cartesian component of *e*_*r*_ is given by *x*_*j*_/*r*. Evaluating the integral between *R*_1_ and *R*_2_ results in

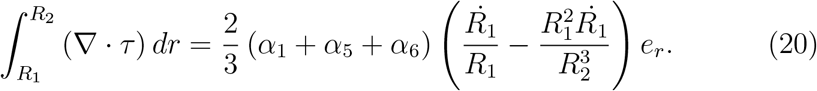

## 4 The elastic energy density for a shelled microbubble

The liquid-crystal shell has both a viscous stress associated with the Leslie viscosities and a stress due to the elastic energy of the liquid crystal. This latter stress will add a further term to equation (9) as calculated below. The following strain energy density function was proposed for a bilipid membrane by De Vita and Stewart [28]

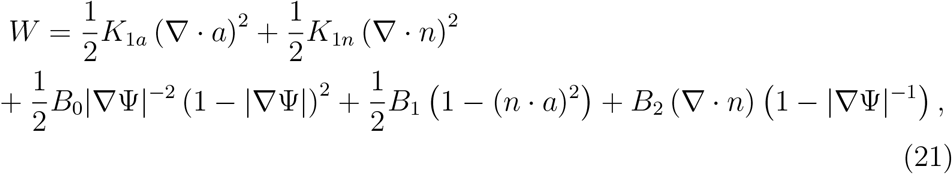

where *K*_1*a*_, *K*_1*n*_, *B*_0_, *B*_1_ and *B*_2_ are material constants, *a* is the unit normal to the layer, Ψ defines the layer structure of a liquid crystal and |∇Ψ|^−1^ represents the current local interlayer distance. The first term on the right hand side of equation (21) refers to the bending energy while the second term represents the splay energy contribution. The *B*_0_ term represents the compression-expansion energy, *B*_1_ is the energy associated with the coupling between *n* and *a*, and *B*_2_ is the term associated with the coupling between the splay and compression-expansion of the layer. It is assumed that the shelled microbubble is a bilipid membrane with a typical thickness of 4nm ([21], p4). Generally |∇Ψ|^−1^ ≠ 1 although for an undistorted liquid-crystal such as planar layers it is useful to define |∇Ψ|^−1^ such that |∇Ψ|^−1^ = 1. There is no contribution to the strain energy density function from the *B*_0_, *B*_1_ and *B*_2_ terms given in equation (21). There are no published values for *K*_1*a*_ but *K*_1*n*_ is known for several types of liquid-crystalline material ([21], p330). We shall make the assumption that *K*_1*a*_ ≈ *K*_1*n*_ such that *K*_1*a*_ = *K*_1*n*_ = *K*_1_. This assumption is based on the experimentally determined values of *K*_1*n*_ for various types of liquid crystals, all of which are very similar in magnitude. Assuming that *n* = *a* then we can conclude that the contribution from the elastic energy density reduces to

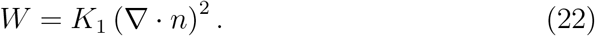

The stress associated with the elastic constant arising from the splay and the bending energies given by *K*_1_(*n*_*i,i*_)^2^ is determined via (*−∂W*/*∂n*_*p,j*_) *n*_*p,i*_ and is represented by *τ*_*elastic*_ ([21],p151) where *W* is given by equation (22). So

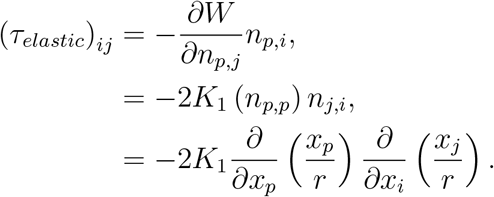

The integral of the divergence of the stress associated with the elastic energy density contributions due to *n* and *a* is

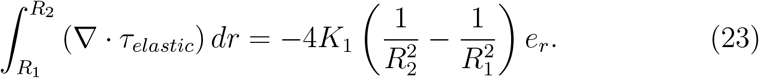

Combining equations (20) and (23) gives the total stress in the shell as

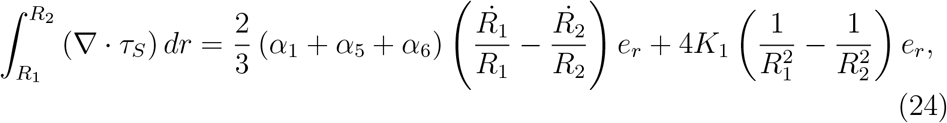

where *τ*_*S*_ represents the total stress in the shell. Substituting equations (7), (8), (18)and (24) into equation (6) gives

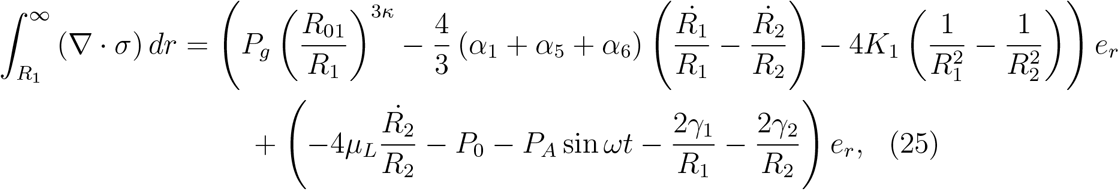

where *R*_01_ is the unperturbed inner radius. To simplify the notation let 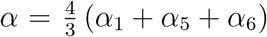. Using equation (23), the stress *S* in its unperturbed state is equal to 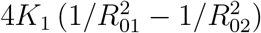. Substituting equation (25) into the right-hand side of equation (1) leads to

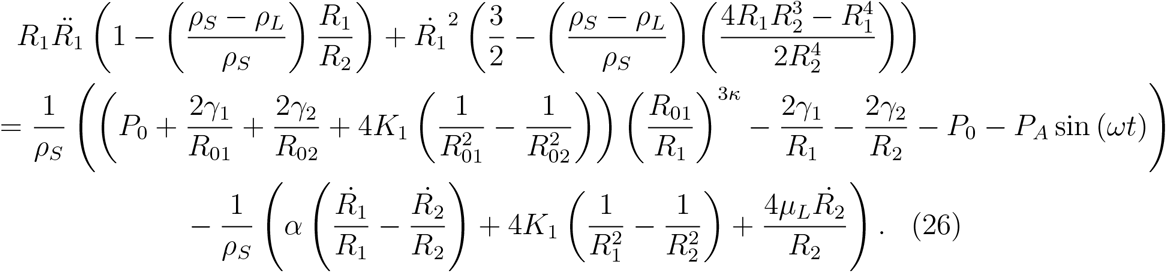

## 5 Linearisation

The technique of linearisation is used to determine the natural frequency and relaxation time for the shelled microbubble whose dynamic behaviour is described by equation (26). The time-dependent perturbations for the inner and outer radii can be written as

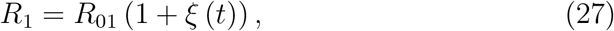

and

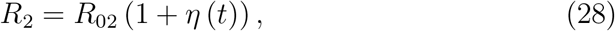

respectively. The shell is incompressible which results in 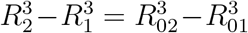. Linearising 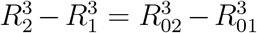 using equations (27) and (28) and assuming that |*ξ*|, |*η*| ≪ 1, results in

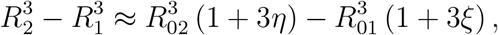

which can be simplified to give

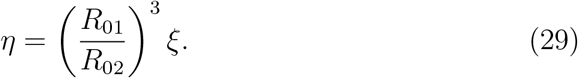

To linearise equation (26) we have to assume that the externally applied forcing pressure *P*_*A*_ is of the same order of magnitude (in some appropriate sense) as |*ξ*| and |*η*|. Then linearising equation (26) leads to

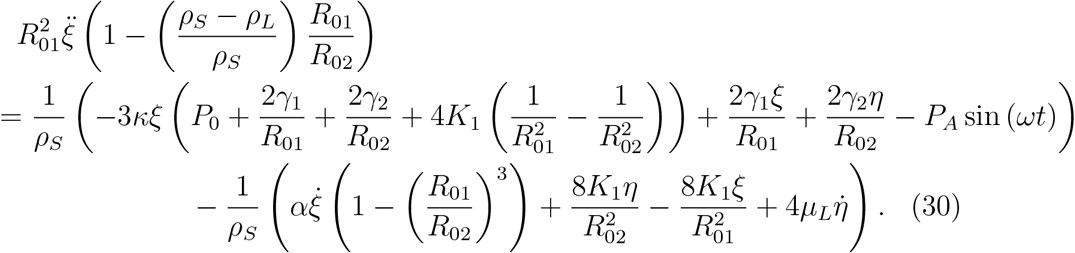

Dividing equation (30) throughout by 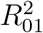 and substituting equation (29) into it gives

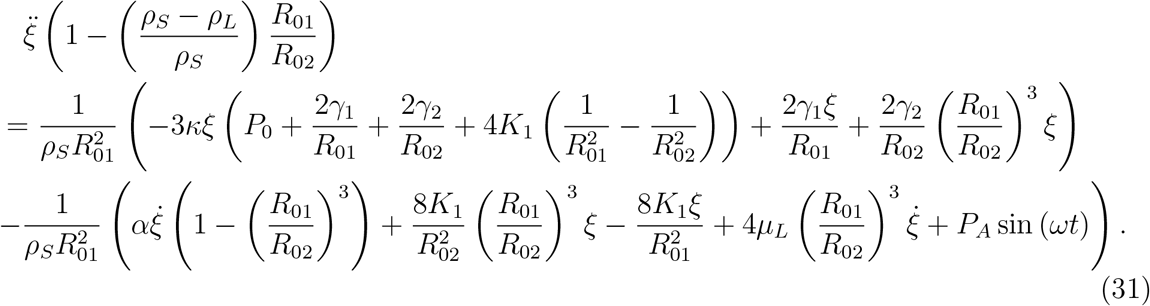

Note that the linearised equation (31) has the form

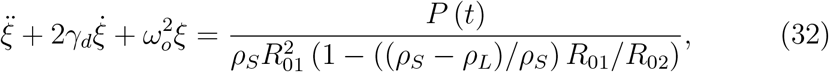

where *γ*_*d*_ represents a damping term and *ω*_*o*_ is the angular natural frequency of the shelled microbubble. The term, *P* (*t*), represents the sinusoidal, external ultrasound signal which forces the shelled microbubble. The damping term is given as

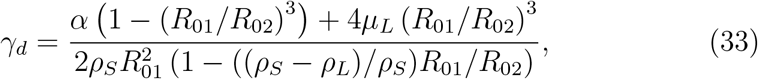

which is related to the relaxation time by *t*_*relax*_ = 1*/γ_d_*. The natural frequency, *f*_*o*_ = *ω_o_/*(2*π*), is given by

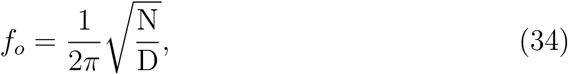

where

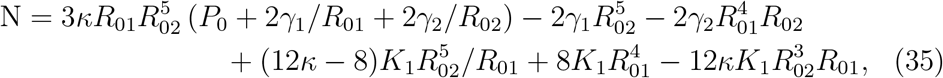

and

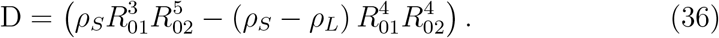

## 6 Results

We will now perform a sensitivity analysis on the damping term *γ*_*d*_ and the natural frequency *f*_0_ given by equations (33) and (34) respectively. We shall consider how *γ*_*d*_ is influenced by changing firstly the Leslie viscosities given by *α*, and then the thickness of the shell given by *R*_02_ − *R*_01_. Also, we shall consider the influence of the interfacial surface tension *γ*_1_ on the natural frequency *f*_0_.

**Figure 1:**
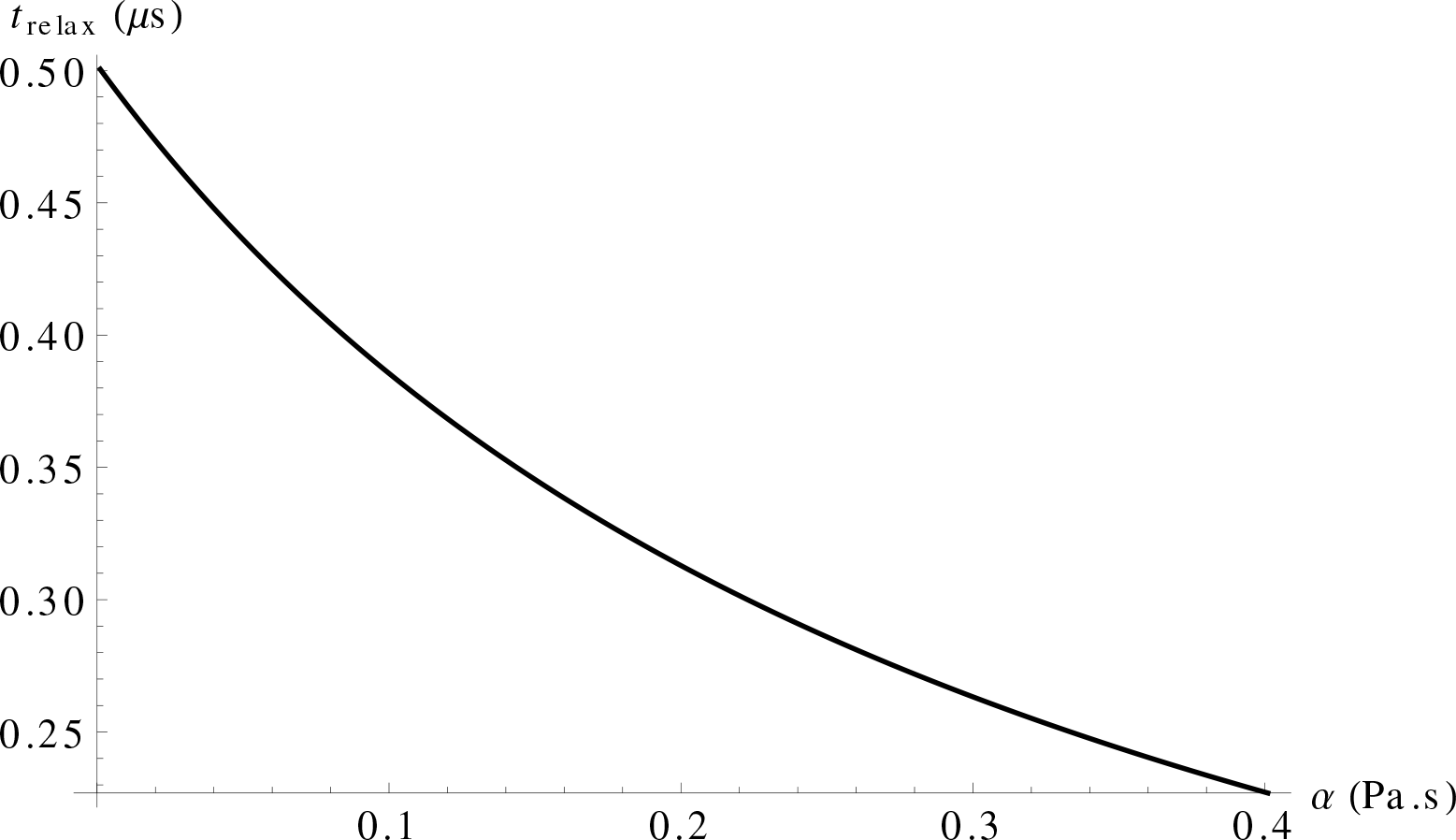
The relaxation time of a shelled microbubble of exterior radius 1*μm* (thickness 4nm) versus the Leslie viscosities given by *α* where 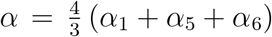. The densities of the liquid-crystalline shell and the surrounding fluid are *ρ*_*S*_ = 1060 kg*m*^−3^ and *ρ*_*L*_ = 1000kg*m*^−3^ respectively ([21], p330). The polytropic index of the gas, the viscosity of the surrounding fluid and the interfacial surface tension and the exterior radius’ surface tension are *κ* = 1.095, *μ*_*L*_ = 10^−3^Pa s, *γ*_1_ = 0.036N*m*^−1^ and *γ*_2_ = 0.072N*m*^−1^ respectively ([21], p330). The graph is constructed using equation (33) and *t*_*relax*_ = 1/*γ*_*d*_.

Figure 1 illustrates the relaxation time’s dependency on the Leslie viscosities where 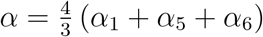. As *α* increases the relaxation time *t*_*relax*_ decreases in a nonlinear manner. This is because a more viscous shell will dampen the oscillatory motion of the shell faster.

**Figure 2:**
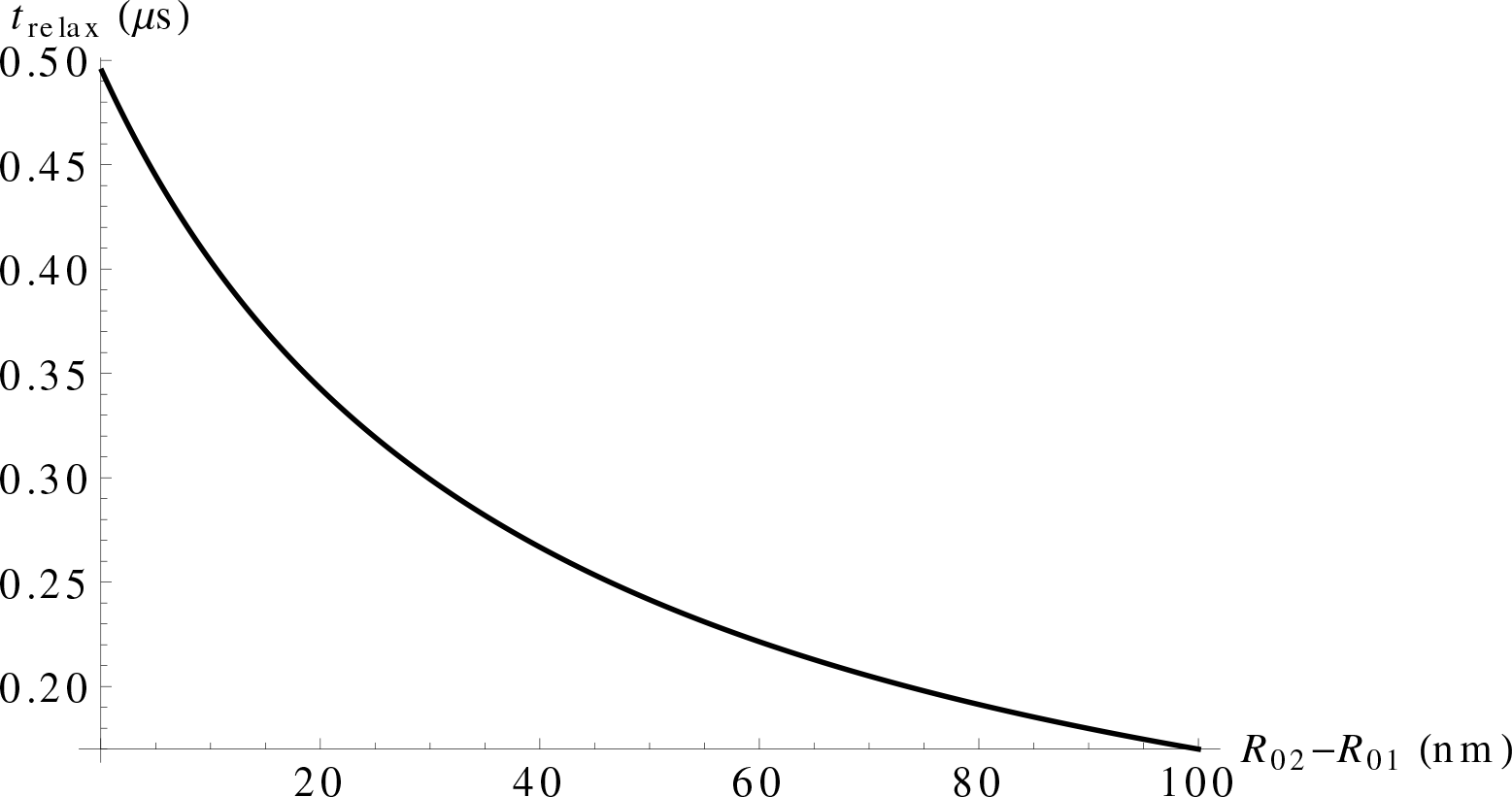
The relaxation time of a shelled microbubble of exterior radius *R*_02_ = 1*μm* versus the thickness of the shell *R*_02_ − *R*_01_. The density of the surrounding fluid and the shell are *ρ*_*L*_ = 1000kg*m*^−3^ and *ρ*_*S*_ = 1060kg*m*^−3^ respectively and the Leslie viscosities give *α* = 0.035Pa s ([21], p330). The polytropic index of the gas, the viscosity of the surrounding fluid and the interfacial surface tension and the exterior radius’ surface tension are *κ* = 1.095, *μ*_*L*_ = 10^−3^Pa s, *γ*_1_ = 0.036N*m*^−1^ and *γ*_2_ = 0.072N*m*^−1^ respectively ([21],p330). The graph is plotted using equation (33) and *t*_*relax*_ = 1/*γ*_*d*_.

Figure 2 shows how the relaxation time decreases nonlinearly as the thickness (*R*_02_ − *R*_01_) of the shell increases. equation (33) highlights the dependency of the damping coefficient and the relaxation time on the radii of the shell. Rewriting equation (33) as

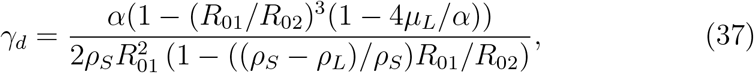

and rearranging for a fixed *α*, *μ*_*L*_, *R*_01_, *ρ*_*S*_ and *ρ*_*L*_ gives

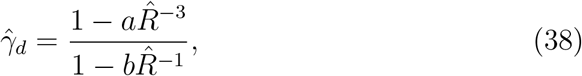

where *a* = 1 − 4*µ*_*L*_/*α*, *b* = (*ρ*_*S*_ − *ρ*_*L*_)/*ρ*_*S*_, 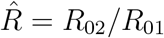 and 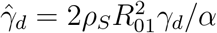. Note that 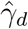 can increase or decrease as a function of 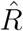 depending on the values of *a* and *b*. Substituting the values of *µ*_*L*_, *α*, *ρ*_*S*_ and *ρ*_*L*_ that are used to construct Figure 2 (see caption) satisfies the condition *a* ≫ *b*. Hence as 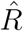 increases and the shell thickens, the damping coefficient *γ*_*d*_ increases which results in a shorter relaxation time (since *t*_relax_ = 1/*γ*_*d*_).

**Figure 3:**
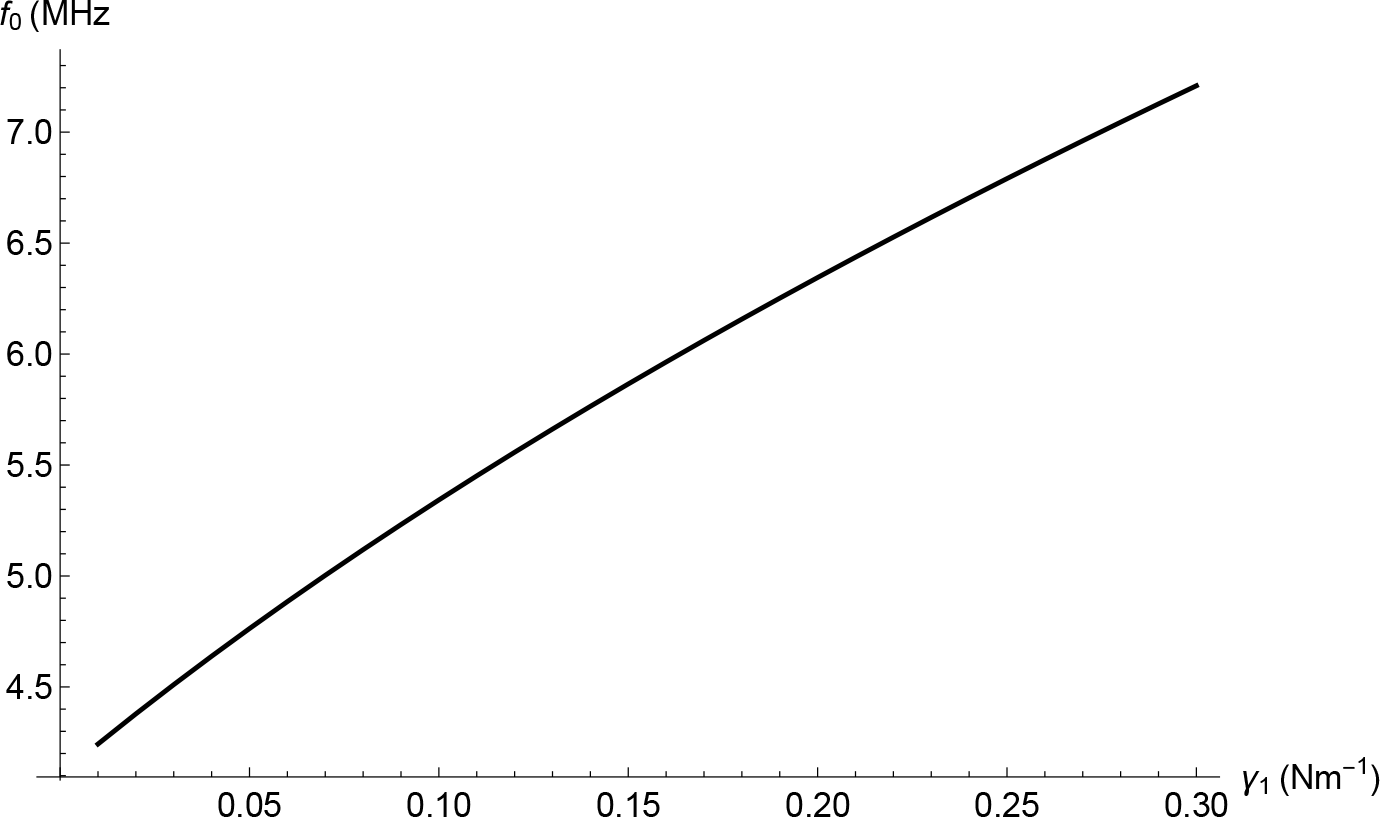
The natural frequency *f*_*o*_ of a shelled microbubble of exterior radius *R*_02_ = 1*μm* and thickness 4nm versus the interfacial surface tension (gas/inner boundary) of the shell denoted by *γ*_1_. The polytropic index of the gas is *κ* = 1.095, the internal gas pressure *P*_0_ = 10^5^Pa [19], the surface tension at the outer shell/liquid interface is *γ*_2_ = 0.072N*m*^−1^ and the elastic constant is given by *K*_1_ = 6 × 10^−12^N ([21], p330). The density of the surrounding fluid and shell are *ρ*_*L*_ = 1000kg*m*^−3^ and *ρ*_*S*_ = 1060kg*m*^−3^ respectively, and the Leslie viscosities results in *α* = 0.035Pa s ([21], p330). The graph is plotted using equations (34), (35) and (36).

Figure 3 illustrates how the natural frequency *f*_*o*_ of the shell increases nonlinearly as the interfacial surface tension *γ*_1_ of the shell increases. This is due to a greater restoring force arising from a larger surface tension acting on the inner surface of the shell.

The known published density of liquid crystals lies between 1020 and 1168 kgm^−3^ ([21], p330), thus varying the density of the shell *ρ*_*S*_ has a negligible effect on both the damping term *γ*_*d*_ and the natural frequency *f*_0_. Similarly varying the elastic constant *K*_1_ over two orders of magnitude has very little effect on the natural frequency *f*_0_. We can compare our theoretically derived expression for the liquid-crystalline shell’s damping term given by equation (33) with the damping term derived by Doinikov and Bouakaz [30] which is for a protein shelled microbubble. Similarly we can compare the natural frequency *f*_0_ given by equation (34) with the natural frequency derived by Doinikov and Bouakaz for a commercial protein shelled microbubble. We shall compare a liquid-crystalline shelled microbubble of thickness *R*_02_ − *R*_01_ and outer equilibrium radius of *R*_02_ = 1*μ*m to an equally sized commercial protein shelled microbubble discussed by Doinikov and Bouakaz [30]. Let us assume that the densities of the liquid-crystalline shell and the surrounding fluid are *ρ*_*S*_ = 1060 kg*m*^−3^ and *ρ*_*L*_ = 1000kg*m*^−3^ respectively ([21], p330). The Leslie viscosity term, the polytropic index of the gas, the viscosity of the surrounding fluid and the interfacial surface tension and the exterior radius’ surface tension are *α* = 0.035Pa s, *κ* = 1.095, *μ*_*L*_ = 10^−3^Pa s, *γ*_1_ = 0.036N*m*^−1^ and *γ*_2_ = 0.072N*m*^−1^ respectively ([21], p330). The damping term for a liquid-crystalline shelled microbubble was *γ*_*d*_ = 2.2 *×* 10^6^s^−1^ compared to *γ*_*d*_ = 3.2*×*10^7^s^−1^ for a commercial protein shelled microbubble. This results in a relaxation time of *t*_relax_ = 4.5 *×* 10^−7^s for a liquid-crystalline shelled microbubble compared to *t*_relax_ = 3.2 *×* 10^−8^s for a commercial protein shelled microbubble. Comparing the natural frequencies *f*_0_ for both types of shells where *P*_0_ = 10^5^Pa gives *f*_0_ = 4.6MHz for a liquid-crystalline shell compared to *f*_0_ = 10.8MHz for a commercial protein shelled microbubble. Note that *f*_0_ = 10.8MHz is the mathematically determined natural frequency for a single protein shelled microbubble. Our study does not consider a uniform solution of microbubbles or a polydisperse solution. Cowley and McGinty have speculated that these significantly different physical characteristics strongly influence the mechanism of sonoporation [17]. Cowley and McGinty have proposed that a liquid-crystalline shelled microbubble enhances the capillary wall shear stress by two orders of magnitude compared to commercial protein shelled microbubbles.

## 7 Conclusion

A modified Rayleigh-Plesset equation has been derived for a shelled microbubble with an incompressible shell composed of a liquid-crystaline material, surrounded by a Newtonian fluid. The model considered the adiabatic gas inside the shelled microbubble, the thin shell’s crystalline material and the surrounding Newtonian fluid. We then linearised the model using time-dependent perturbation theory and determined expressions for the relaxation time and the natural frequency of the shelled microbubble. We performed a sensitivity analysis, considering how the various material parameters of the shell influenced both the relaxation time and the natural frequency of the shelled microbubble. The relaxation time exhibited a dependency on both the thickness of the shell and the Leslie viscosities (which are depen-dent on the type of liquid-crystalline material that the shell is made from). We discovered that the relaxation time decreased nonlinearly as the Leslie viscosities of the shell increased. Similarly the relaxation time decreased nonlinearly as the thickness of the shell increased. However, the natural frequency of the shelled microbubble depended primarily on the interfacial surface tension of the liquid-crystalline shell. Our sensitivity analysis on the natural frequency showed that the natural frequency increased nonlinearly as the interfacial surface tension increased. Up until now there has been no published experimental data for liquid-crystalline shelled microbubbles. Using the values given by Doinikov and Bouakaz for commercial shelled microbubbles, we have discovered that the damping term *γ*_*d*_ for commercial microbubbles is approximately 10 times larger than the damping term for a liquid-crystalline shelled microbubble. This implies that current commercial shelled microbubbles have a relaxation time that is approximately 10 times shorter. We have also discovered that the natural frequency of a liquid-crystalline shelled microbubble is approximately 1/2 that of a commercial shelled microbubble. There are two novel contributions in this article. We have derived for the first time a modified Rayleigh-Plesset equation for a shelled microbubble whose shell is composed of a liquid-crystalline material. No previous study has considered such an alternative and unique approach. We have given qualitative insight into how the material parameters such as the Leslie viscosities, the thickness of the shell, and the interfacial surface tension of the shell influence the shelled microbubble’s relaxation time and natural frequency. Such modelling may aid soft matter scientists’ understanding of UCA localised drug delivery and gene therapy specifically in the treatment of cancer.

Future research will focus on the technique of sonoporation which involves using the shelled microbubbles in conjunction with an external ultrasound signal to temporarily enhance the porosity of the capillary walls. This temporary enhancement of the walls is a consequence of wall shear stress and is due to several mechanisms [30]. One such mechanism is acoustic microstreaming which we intend to model using our liquid-crystalline shelled microbubble model. It has been proposed by Doinikov and Bouakaz that both the damping term and the natural frequency of the shell have a significant influence on the magnitude of the wall shear stress. We will compare and contrast the wall shear stress generated by a solution of liquid-crystalline shelled microbubbles acting on a viscoelastic capillary wall to that generated by a solution of commercial contrast agents such as Sonovue.

We accept that experimental data is required in order to validate the findings of our mathematical model. It is the authors’ hope that this journal article instigates future experimental work.

## 8 Acknowledgements

The authors gratefully acknowledge the support given by the UK Engineering and Physical Sciences Research Council via a Doctoral Training Grant [grant number EP/L505080/1].

## References

[1] P. Narayan and M.A. Wheatley. Preparation and characterization of hollow microcapsules for use as ultrasound contrast agents. Polymer Engineering and Science, 39:2242–2255, 1999.

[2] M. Bazan Peregrino, B. Rifai, R.C. Carlisle, J. Choi, C.D. Arvanitis, L.W. Seymour, and C.C. Coussios. Cavitation-enhanced delivery of a replicating oncolytic adenovirus to tumors using focused ultrasound. Journal of Controlled Release, 169:40–47, 2013.

[3] D. Gourevich, A. Volovick, O. Dogadkin, L. Wang, H. Mulvana, Y. Medan, A. Melzer, and S. Cochran. In vitro investigation of the individual contributions of ultrasound-induced stable and inertial cavitation in targeted drug delivery. Ultrasound in Medicine & Biology, 41:1853–1864, 2015.

[4] J.M. Escoffre, C. Mannaris, B. Geers, A. Novell, I. Lentacker, M. Averkion, and A. Bouakaz. Doxirubicin liposome-loaded microbubbles for contrast imaging and ultrasound triggered drug delivery. IEEE Transactions on Ultrasonics, Ferroelectrics and Frequency Control, 60:78–87, 2013.

[5] C.H. Fan, C.Y. Ting, H.L. Liu, C.Y. Huang, H.Y. Hsieh, T.C. Yen, K.C. Wei, and C.K. Yeh. Antiangiogenic-targeting drug-loaded microbubbles combined with focused ultrasound for glioma treatment. Biomaterials, 34:2142–2155, 2013.

[6] F. Yan, L. Li, Z. Deng, Q. Jin, J. Chen, W. Yang, C.K. Yeh, J. Wu, R. Shandas, X. Liu, and H. Zheng. Paclitaxel-liposome-microbubble complexes as ultrasound-triggered therapeutic drug delivery carriers. Journal of Controlled Release, 166:246–255, 2013.

[7] C. Niu, Z. Wang, G. Lu, T.M. Krupka, Y. Sun, Y. You, W. Song, H. Ran, P. Li, and Y. Zheng. Doxorubicin loaded superparamagnetic PLGA-iron oxide multifunctional microbubbles for dual-mode US/MR imaging and therapy of metastasis in lymph nodes. Biomaterials, 34:2307–2317, 2013.

[8] A. Delalande, C. Leduc, P. Midoux, M. Postema, and C. Pichon. Efficient gene delivery by sonoporation is associated with microbubble entry into cells and the clathrin-dependent endocytosis pathway. Ultrasound in Medicine & Biology, 41:1913–1926, 2015.

[9] C. McEwan, J. Owen, E. Stride, C. Fowley, H. Nesbitt, D. Cochrane, C.C. Coussios, M. Borden, N. Nomikou, A.P. McHale, and J.F. Callan. Oxygen carrying microbubbles for enhanced sonodynamic therapy of hypoxic tumours. Journal of Controlled Release, 203:51–56, 2015.

[10] E. Stride and N. Saffari. Microbubble ultrasound contrast agents: a review. Proceedings of the Institution of Mechanical Engineers Part H, 217:429–447, 2003.

[11] D. Cosgrove. Ultrasound contrast agents: An overview. European Journal of Radiology, 60:324–330, 2006.

[12] E. Stride and C.C. Coussios. Cavitation and contrast: The use of bubbles in ultrasound imaging and therapy. Proceedings of the Institute of Mechanical Engineers Part H, 224:171–191, 2010.

[13] J. McLaughlan, N. Ingram, P.R. Smith, S. Harput, P.L. Coletta, S. Evans, and S. Freear. Increasing the sonoporation efficiency of targeted polydisperse microbubble populations using chirp excitation. IEEE Transactions on Ultrasonics, Ferroelectrics and Frequency Control, 60:2511–2520, 2013.

[14] M.J.K. Blomley, J.C. Cooke, E.C. Unger, M.J. Monaghan, and D.O. Cosgrove. Microbubble contrast agents: a new era in ultrasound. British Medical Journal, 322:1222–1225, 2001.

[15] I. Lentacker, S.C. De Smedt, and N.N. Sanders. Drug loaded microbubble design for ultrasound triggered delivery. Soft Matter, 5:2161–2170, 2009.

[16] H. Dewitte, K. Vanderperren, H. Haers, E. Stock, L. Duchateau, M. Hesta, J.H. Saunders, S.C. De Smedt, and I. Lentacker. Theranostic mRNA-loaded microbubbles in the lymphatics of dogs:implications for drug delivery. Theranostics, 5:97–109, 2015.

[17] J. Cowley and S. McGinty. A mathematical model of sonoporation using a liquid-crystalline shelled microbubble. Ultrasonics, https://doi.org/10.1016/j.ultras.2019.01.004.

[18] C.C. Church. The effects of an elastic solid surface layer on the radial pulsations of gas bubbles. Journal Acoustical Society of America, 97:1510–1521, 1995.

[19] P. Marmottant, S. Van der Meer, M. Emmer, M. Versluis, N. de Jong, S. Hilgenfeldt, and D. Lohse. A model for large amplitude oscillations of coated bubbles accounting for buckling and rupture. Journal Acoustical Society of America, 118:3499–3505, 2005.

[20] A.A Doinikov and A. Bouakaz. Review of shell models for contrast agent microbubbles. IEEE Transactions on Ultrasonics, Ferroelectrics and Frequency Control, 58:981–993, 2011.

[21] I.W. Stewart. The Static and Dynamic Continuum Theory of Liquid Crystals. Taylor and Francis, London, 2004.

[22] A.A. Doinikov and P.A. Dayton. Maxwell rheological model for lipid-shelled ultrasound microbubble contrast agents. Journal Acoustical Society of America, 121:3331–3340, 2007.

[23] A.A. Doinikov, J.F. Haac, and P.A. Dayton. Modeling of nonlinear viscous stress in encapsulating shells of lipid-coated contrast agent microbubbles. Ultrasonics, 49:269–275, 2009.

[24] S. Paul, A. Katiyar, K. Sarkar, D. Chatterjee, W.T. Shi, and F. Forsberg. Material characterization of the encapsulation of an ultrasound contrast microbubble and its subharmonic response: Strain-softening interfacial elasticity model. Journal Acoustical Society of America, 127:3846–3857, 2010.

[25] C. Brennen. Cavitation and Bubble Dynamics. Oxford University Press, New York, 1995.

[26] D.J. Acheson. Elementary Fluid Dynamics. Oxford University Press, New York, 1990.

[27] F.M. Leslie. Continuum theory for nematic liquid crystals. Continuum Mechanics and Thermodynamics, 4:167–175, 1992.

[28] R. De Vita and I.W. Stewart. Energetics of lipid bilayers with applications to deformations induced by inclusions. Soft Matter, 9:2056–2068, 2013.

[29] I.W. Stewart. Dynamic theory for smectic A liquid crystals. Continuum Mechanics and Thermodynamics, 18:343–360, 2007.

[30] A.A. Doinikov and A. Bouakaz. Theoretical investigation of shear stress generated by a contrast microbubble on the cell membrane as a mechanism for sonoporation. Journal Acoustical Society of America, 128:11–19, 2010.

